# Abnormal cell sorting underlies the unique X-linked inheritance of *PCDH19* Epilepsy

**DOI:** 10.1101/178822

**Authors:** Daniel T. Pederick, Kay L. Richards, Sandra G. Piltz, Simone A. Mandelstam, Russell C. Dale, Ingrid E. Scheffer, Jozef Gecz, Steve Petrou, James N. Hughes, Paul Q. Thomas

**Affiliations:** School of Biological Sciences and Robinson Research Institute, The University of Adelaide, Adelaide, South Australia 5005, Australia; Florey Institute of Neuroscience and Mental Health, The University of Melbourne, Melbourne, Victoria 3010, Australia; Department of Paediatrics, The University of Melbourne, Melbourne, Victoria 3010, Australia; Department of Radiology, The University of Melbourne, Melbourne, Victoria 3010, Australia; Department of Medical Imaging, Royal Children’s Hospital, Florey Neurosciences Institute, Parkville, Victoria 3052, Australia; Institute for Neuroscience and Muscle Research, University of Sydney; University of Melbourne, Austin Health and Royal Children’s Hospital, Victoria, 3084, Australia; School of Medicine, The University of Adelaide, Adelaide, South Australia 5005, Australia; South Australian Health and Medical Research Institute, Adelaide, South Australia 5000, Australia

**Keywords:** Adhesion molecules, Protocadherins, Protocadherin 19, cell-cell adhesion code, epilepsy, cell sorting, cortical development, *PCDH19*-GCE

## Abstract

X-linked diseases typically exhibit more severe phenotypes in males than females. In contrast, *Protocadherin 19* (*PCDH19*) mutations cause epilepsy in heterozygous females but spare hemizygous males. The cellular mechanism responsible for this unique pattern of X-linked inheritance is unknown. We show that PCDH19 contributes to highly specific combinatorial adhesion codes such that mosaic expression of *Pcdh19* in heterozygous female mice leads to striking sorting between WT PCDH19- and null PCDH19-expressing cells in the developing cortex, correlating with altered network activity. Complete deletion of PCDH19 in heterozygous mice abolishes abnormal cell sorting and restores normal network activity. Furthermore, we identify variable cortical malformations in PCDH19 epilepsy patients. Our results highlight the role of PCDH19 in determining specific adhesion codes during cortical development and how disruption of these codes is associated with the unique X-linked inheritance of *PCDH19* epilepsy.

## Introduction

Protocadherins (PCDHs) are the largest family (∼70 genes) of cell-cell adhesion molecules and have important roles in many neurobiological processes including axon guidance/sorting, neurite self-avoidance and synaptogenesis (Garrett and Weiner, 2009; Lefebvre et al., 2012; Uemura et al., 2007). The majority of PCDHs are positioned within three genomic clusters termed α, β and γ. Clustered PCDHs are stochastically expressed in neurons which is thought to be essential for generating individual neuron identity (Rubinstein et al., 2015; Schreiner and Weiner, 2010; Thu et al., 2014). The remaining family members are termed non-clustered (NC) PCDHs. Genes encoding NC PCDHs are scattered throughout the genome and display unique and overlapping expression patterns in different neuronal populations during central nervous system (CNS) development and in the mature brain (Kim et al., 2007; Krishna-K, 2009).

Mutations in NC PCDH family members has been associated with a variety of neurological disorders including epilepsy, autism and schizophrenia (Bray et al., 2002; International League Against Epilepsy Consortium on Complex Epilepsies., 2014; Ishizuka et al., 2016; Morrow et al., 2008). Notably, mutations in Protocadherin 19 (*PCDH19)* cause *PCDH19* Girls Clustering Epilepsy (*PCDH19*-GCE) which is reported to be the second most common cause of monogenic epilepsy (Dibbens et al., 2008; Duszyc et al., 2014). *PCDH19*-GCE is an X-linked disorder with a unique pattern of inheritance whereby heterozygous females are affected while hemizygous males are spared (Dibbens et al., 2008; Ryan et al., 1997). It has been proposed that the co-existence of wild type (WT) and mutant *PCDH19* neurons that arise through random X-inactivation underpins the unique inheritance (Depienne et al., 2009; Dibbens et al., 2008). This hypothesis is supported by the existence of affected males who are mosaic carriers of somatic *PCDH19* mutations (Depienne et al., 2009). However, it is currently unclear what processes are regulated by PCDH19 in the brain and how these are disrupted by mosaic expression but not by the complete absence of functional protein.

PCDH19 has been shown to function as a homotypic cell adhesion molecule *in vitro* (Tai et al., 2010) and recently published data indicate that PCDH19 has a virtually identical structure to members of the closely-related clustered PCDH family (Cooper et al., 2016). Interestingly, clustered PCDHs can form heterotypic *cis* interactions which generate highly specific adhesion codes that are likely to play a role in vertebrate neuronal self-avoidance (Rubinstein et al., 2015; Schreiner and Weiner, 2010; Thu et al., 2014). Interestingly, NC PCDHs display overlapping expression *in vivo* (Kim et al., 2007; Krishna-K, 2009), suggesting they may also contribute to adhesion codes in a combinatorial manner. However, it remains unclear whether PCDH19 can form heterotypic *cis* interactions with other NC PCDHs and contribute to combinatorial cell-cell adhesion codes.

Here, we sought to address these questions by performing complementary *in vitro* and *in vivo* experiments to investigate the combinatorial activity of PCDH19 and the molecular mechanism underlying the unique X-linked inheritance pattern of *PCDH19*-GCE. Our findings show that PCDH19 contributes to specific combinatorial adhesion codes such that mosaic expression of PCDH19 *in vivo* disrupts cell adhesion specificity resulting in abnormal sorting of neuroprogenitor cells in the developing cortex.

## Results

### PCDH19 Forms Heterotypic *cis* Interactions with NC PCDHs that Generate Highly Specific Binding Affinities

To investigate the adhesion activity of PCDH19 in combination with other NC PCDHs we performed mixing experiments using K562 cells which ordinarily do not aggregate during culture due to a lack of endogenous PCDHs and classical cadherins (Ozawa and Kemler, 1998; Schreiner and Weiner, 2010). PCDH17 and PCDH10 were selected for these experiments due to their ability to interact in *cis* with PCDH19 when co-expressed in the same cell in a co-immunoprecipitation assay (Figure 1A) and overlapping expression with *Pcdh19* in the embryonic mammalian brain (Kim et al., 2007). K562 cells transfected with different fluorescently-labelled PCDH combinations were mixed in a 1:1 ratio and quantitatively assessed for the presence/absence of sorting between the two populations (Figure 1B). When both cell populations expressed the same combination of PCDHs random mixing was observed (Figure 1C). In contrast, populations expressing different PCDH combinations exhibited significant cell sorting indicating that cells with matching PCDH profiles preferentially adhere (Figure 1C). Notably, significant sorting occurred between populations that differed by a single PCDH. We next assessed the impact of *PCDH19*-GCE missense mutations on combinatorial PCDH adhesion. We first individually expressed PCDH19 containing one of three commonly occurring missense mutations in K562 cells and found that each lacked adhesive function (Figure S1), consistent with previously reported data for other missense mutations (Cooper et al., 2016). When PCDH10 was co-expressed with either WT PCDH19 or mutant PCDH19.N340S, significant sorting occurred between the two populations (Figure 1C). Taken together, these data suggest that PCDH19 forms heterotypic *cis* interactions with NC PCDHs and contributes to combinatorial adhesion codes that are sensitive to single PCDH differences.

**Figure 1.**
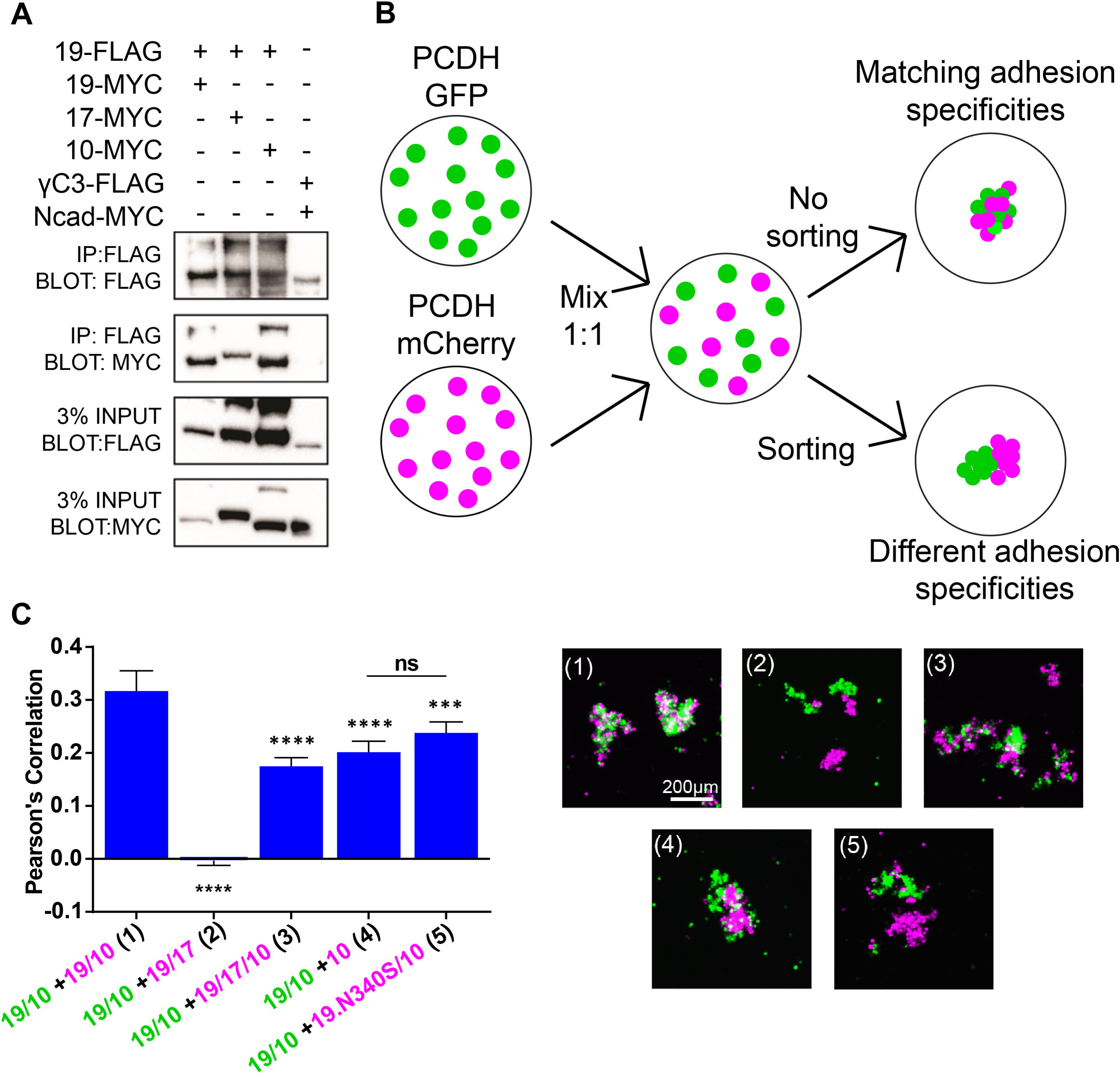
δ2 NC PCDHs form combinatorial codes which dictate adhesion specificity. (A) Co-immunoprecipitation of K562 cell lysates expressing NC PCDH combinations indicated by the + symbol. Lysates were pulled-down with FLAG Ab and subsequent blotting was performed with MYC Ab (*N*=3 experiments). Cells transfected with PCDHγC3-FLAG and Ncad-MYC served as a negative control for cis-interaction as previously reported (Thu et al., 2014). (B) Schematic diagram describing mixing assays performed to assess adhesion specificity. (C) Quantification of the degree of mixing using Pearson’s correlation coefficient. Expression constructs are indicated by 19 (PCDH19), 17 (PCDH17), 10 (PCDH10) and 19. N340S (PCDH19 N340S missense mutation) (\*\*\**P* < 0.001, \*\*\*\**P* < 0.0001, ns=not significant, one-way analysis of variance (ANOVA) with Tukey’s multiple comparison test against 19/10 +19/10 unless otherwise indicated). Representative images of quantified K562 mixing assays. See Table S1 for additional information including *n* for each experiment.

### Mosaic Expression of *Pcdh19 in vivo* Leads to Altered Network Brain Activity Correlating with Abnormal Cell Sorting in the Developing Cortex

Next, we sought to investigate how perturbation of these adhesion codes would manifest in the developing mammalian brain where complex arrays of adhesion molecules direct morphogenesis. For these experiments we used mice carrying a null allele (*Pcdh19*^*β-Geo*^) (Pederick et al., 2016) to generate female heterozygous mice with mosaic expression of *Pcdh19.* Given the seizures and elevated neural activity of *PCDH19*-GCE affected females (Higurashi et al., 2013; Scheffer et al., 2008), we initially performed electrocorticogram (ECoG) analysis on young adult mice to investigate if a phenotype exists in heterozygous mice that does not manifest in homozygote animals. ECoG traces from *Pcdh19^+/^*^*β-Geo*^ (*+/β-Geo*) postnatal day 42 (P42) mice showed a consistent increase in amplitude compared to *Pcdh19^+/+^* (*+/+*) and *Pcdh19*^*β-Geo/β-Geo*^ (*β-Geo/β-Geo*) animals, which were themselves indistinguishable (Figure 2A). Strikingly, ECoG signatures for *+/β-Geo* mice showed a significant increase in mean number of spike-wave discharge (SWD) events per hour and event duration compared with *+/*+ mice (Figure 2B,C; Figure S2). In contrast, the mean number of SWDs per hour and the mean duration of a SWD event for *β-Geo/β-Geo* animals was not significantly different to *+/*+ mice (Figure 2B,C; Figure S2). This indicates that mosaic expression of *Pcdh19* in mice results in altered brain network activity. In contrast, this phenotype is not present in mice completely lacking PCDH19, consistent with the unique X-linked inheritance of *PCDH19*-GCE in humans.

**Figure 2.**
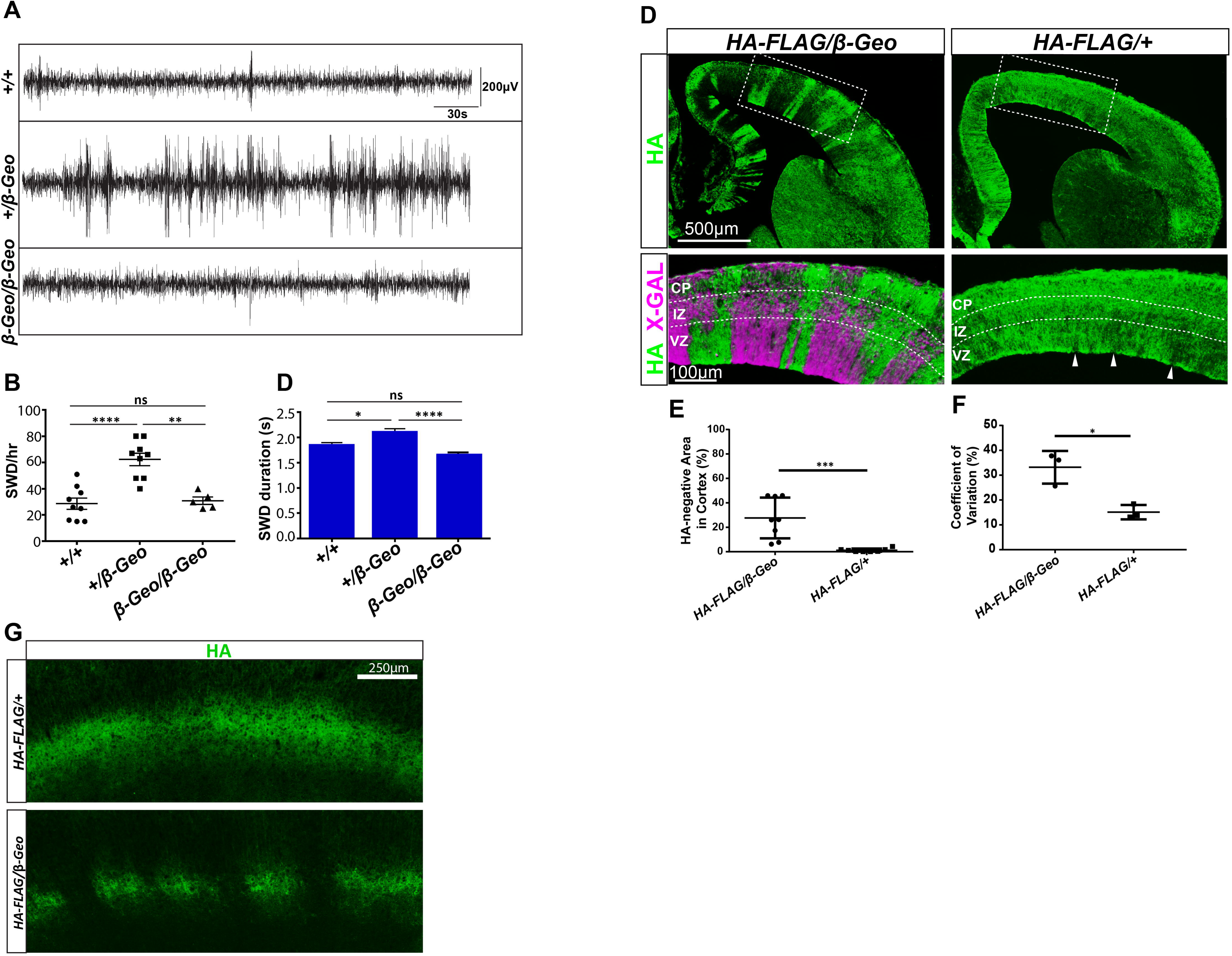
Mosaic expression of *Pcdh19* causes altered network brain activity and abnormal cell sorting between PCDH19 positive and PCDH19 negative cells in the developing cortex. Representative traces from ECoG recordings of *+/+*, *+/β-Geo* and *β-Geo/β-Geo* P42 mice. Quantification of SWD/hr of *+/+*, *+/β-Geo* and *β-Geo/β-Geo* P42 mice (\*\*\*\**P* < 0.0001, \*\**P* < 0.01, ns=not significant, student’s two-tailed, unpaired *t* test *P* values). (C) Quantification of mean SWD duration of *+/+ +/β-Geo* and *β-Geo/β-Geo* P42 mice (\*\*\*\**P* < 0.0001, \**P* < 0.05, ns=not significant, two-way ANOVA with Tukey’s multiple comparisons test). (D) Representative HA and X-gal immunostaining of 14.5 dpc *HA-FLAG/+* and *HA-FLAG/β-Geo* brains. (E) Quantification of HA-negative regions in the cortex of 14.5 dpc *HA-FLAG/β-Geo* and *HA-FLAG/+* and brains (\*\**P* < 0.01, unpaired *t* test). (F) Quantification of variability of HA-immunostaining intensity throughout the cortex (\**P* < 0.05, unpaired *t* test). (G) Representative HA immunostaining of P7 *HA-FLAG/+* and *HA-FLAG/β-Geo* brains.Abbreviations VZ, ventricular zone; IZ, intermediate zone; CP, cortical plate. Also see Table S1.

To determine the impact of mosaic PCDH19 expression at the cellular level *in vivo* we required reporter alleles for *Pcdh19*-expressing WT and null cells. To label *Pcdh19-*expressing null cells, we used the *Pcdh19*^*β-Geo*^ *β-galactosidase* knock-in reporter allele, which we had previously validated using X-Gal staining (Pederick et al., 2016). To enable unequivocal identification of WT PCDH19-expressing cells, we employed CRISPR/Cas9 genome editing to insert an HA-FLAG epitope sequence at the C-terminus of the PCDH19 ORF (Figure S3A). Successful insertion was validated via PCR, sequencing, western blot and HA immunohistochemistry (Figure S3B, C and D, respectively). Consistent with previously published mRNA *in situ* hybridization data (Dibbens et al., 2008; Gaitan and Bouchard, 2006; Pederick et al., 2016), we detected PCDH19 expression in many brain regions, including prominent expression in the developing cortex that became more restricted in the postnatal brain (Figure S3D and E; Figure S4).

Simultaneous labelling of PCDH19 positive (HA) and negative (X-gal) cells in *Pcdh19^HA-FLAG/^*^*β-Geo*^ (*HA-FLAG/β-Geo*) brains enabled us to investigate the phenotype resulting from disruption of PCDH19-depedent cell adhesion codes. HA immunostaining of *HA-FLAG/β-Geo* brains revealed a striking alternating pattern of PCDH19-positive and PCDH19-negative cells that extended from the ventricular zone to the cortical plate. (Figure 2D, left). X-gal staining of *HA-FLAG/β-Geo* brains revealed a complementary and non-overlapping pattern to HA immunostaining (Figure 2D, left), suggesting this pattern is due to segregation of PCDH19-positive and PCDH19-null cells. To confirm that this pattern did not arise due to random X-inactivation and subsequent clonal expansion, we examined HA-staining in “wild type” *Pcdh19^HA-FLAG/+^* (*HA-FLAG/+*) control embryos (Figure 2D, right). This allowed us to identify the pattern of PCDH19-positive HA labelled cells in the presence of PCDH19-positive unlabelled cells. Within the cortex, X-inactivation manifested as small interspersed patches of HA-positive and HA-negative staining along the ventricle and overlying neural progenitors in the ventricular zone, with only subtle variations in HA staining within the cortical plate (Figure 2D, right). The significantly increased percentage of HA-negative regions in *HA-FLAG/β-Geo* embryos compared with *HA-FLAG/+* controls confirms that PCDH19-positive and PCDH19-null cells coalesced into distinct groups (Figure 2E). We also noted significantly increased variation of HA-immunostaining across different *HA-FLAG/β-Geo* animals, suggesting differences in abnormal cell sorting phenotypes are due to unique patterns of X-inactivation in each embryo (Figure 2F; Figure S5; Figure S6). The striking segregation occurring cortical development also persisted postnatally amongst cells that continued to express PCDH19 (Figure 2G). Consistent with an active role in cell sorting, PCDH19 was present at interfaces between PCDH19 expressing neuroprogenitor cells but not between PCDH19-positive and negative cells, supporting its role as a homotypic cell-cell adhesion molecule *in vivo* (Figure S7). Cell body redistribution was independently confirmed by staining for nuclear-localised SOX3 in *Pcdh19^+/^*^*β-Geo*^; *Sox3^+/-^* trans-heterozygous mice (Figure S8). Notably, cell segregation was not detected at the inception of PCDH19 expression at 9.5dpc but was clearly present at 10.5dpc, indicating rapid onset of the abnormal cell sorting phenotype (Figure S9). Taken together, these data suggest that PCDH19 functions to specify cell-cell interactions and that mosaic expression results in distinct cell populations with incompatible adhesion codes which abnormally segregate during cortical development.

### Uniform Deletion of *Pcdh19* Prevents Abnormal Cell Sorting

The absence of pathology in hemizygous males and the normal brain activity of mice completely lacking functional PCDH19 (Figure 2A,B and C) suggests that the removal of differential adhesion codes dictated by mosaic PCDH19 expression will restore normal cell sorting during cortical development. To test this hypothesis, we developed a method to assess *in vivo* cell sorting by visualising *Pcdh19* allele-specific expression in functionally null PCDH19 female mice. Using CRISPR/Cas9 genome editing, we deleted the *Pcdh19^HA-FLAG^* allele in *HA-FLAG/β-Geo* zygotes to create *DEL/β-Geo* null embryos (Figure 3A). Deletion of exon 1 and lack of functional PCDH19 was validated by sequencing and HA immunostaining (Figure 3B and data not shown). X-gal staining of *DEL/β-Geo* embryos at 14 days post embryo transfer demonstrated that the two populations of null cells readily intermix, thereby rescuing the abnormal cell sorting phenotype observed in *HA-FLAG/β-Geo* embryos (Figure 3B). Quantification of X-gal staining variation confirmed that the abnormal cell sorting was rescued in *DEL/β-Geo* cortices (Figure 3C; Figure S5). This provides further evidence that the differential adhesion codes of PCDH19-positive and PCDH19-negative cells leads to abnormal cell sorting. Importantly, the lack of abnormal cell sorting in *DEL/β-Geo* null embryos provides a clear cellular phenotype that correlates with and likely explains the unique X-linked inheritance of *PCDH19* epilepsy.

**Figure 3.**
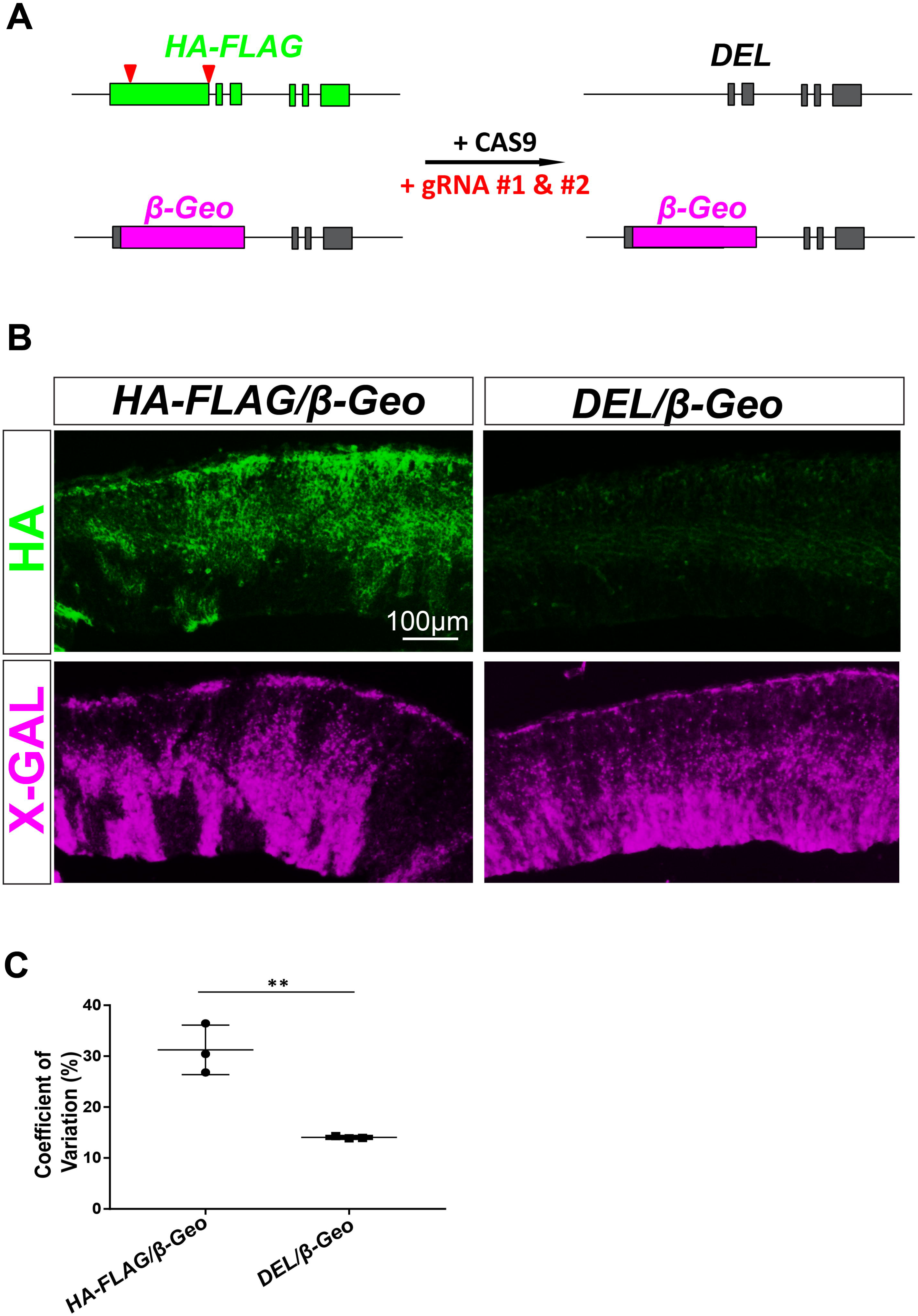
Deletion of *Pcdh19* in *HA-FLAG/β-Geo* mice abolishes abnormal cell sorting. (A) Schematic describing CRISPR/Cas9-induced conversion of *HA-FLAG/β-Geo* embryos to DEL/*β*-Geo. Two gRNAs targeting exon1 flanking sequences (red arrows) were injected into *HA-FLAG/β-Geo* zygotes in combination with CAS9 protein. Zygotes were transferred into pseudopregnant female mice and harvested 14 days post transfer. (B) HA and X-gal immunostaining of *HA-FLAG/β-Geo* and *DEL/β-Geo* 14 day post transfer embryonic brains). (C) Quantification of variability of X-gal staining intensity throughout the ventricle of *HA-FLAG/β-Geo* and *HA-FLAG/+* brains (\*\**P* < 0.01, unpaired *t* test).

### Variable Cortical Folding Abnormalities are Observed in *PCDH19*-GCE Patients

Although structural brain abnormalities have not previously been described in *PCDH19-GCE* affected individuals (Guerrini et al., 2014) we hypothesized that abnormal cell sorting caused by mosaic expression of PCDH19 during human cortical development could result in morphological defects due to the prolonged expansion of neuroprogenitors and extensive sulcation relative to mice (Sun and Hevner, 2014). Using data from the Allen Brain Atlas we confirmed *PCDH19* is highly expressed in human embryonic CNS including the cortex during the key neurogenic period of 8-16 weeks post conception (Miller et al., 2014; Semple et al., 2013) (Figure S10A). *PCDH19* expression is reduced postnatally, but is still detectable, with the brain remaining the predominant site of expression in the adult (Figure S10B). We then reviewed MRI images from a cohort of *PCDH19*-GCE girls. We identified abnormal cortical sulcation in four patients with common causative mutations in *PCDH19* (p.N340S, p.S671X and p.Y366LfsX10; Figure 4; Figure S11; Table S2) (Scheffer et al., 2008). The patients presented with variably positioned cortical defects that included bottom of the sulcus dysplasias, abnormal cortical folding, cortical thickening and blurring of the grey/white junction. These data suggest that subtle brain abnormalities are a feature of *PCDH19-*GCE. While it is not known how abnormal cell sorting could generate these defects, their variability is consistent with the random nature of X-inactivation and phenotypes observed in *PCDH19*-GCE patients.

**Figure 4.**
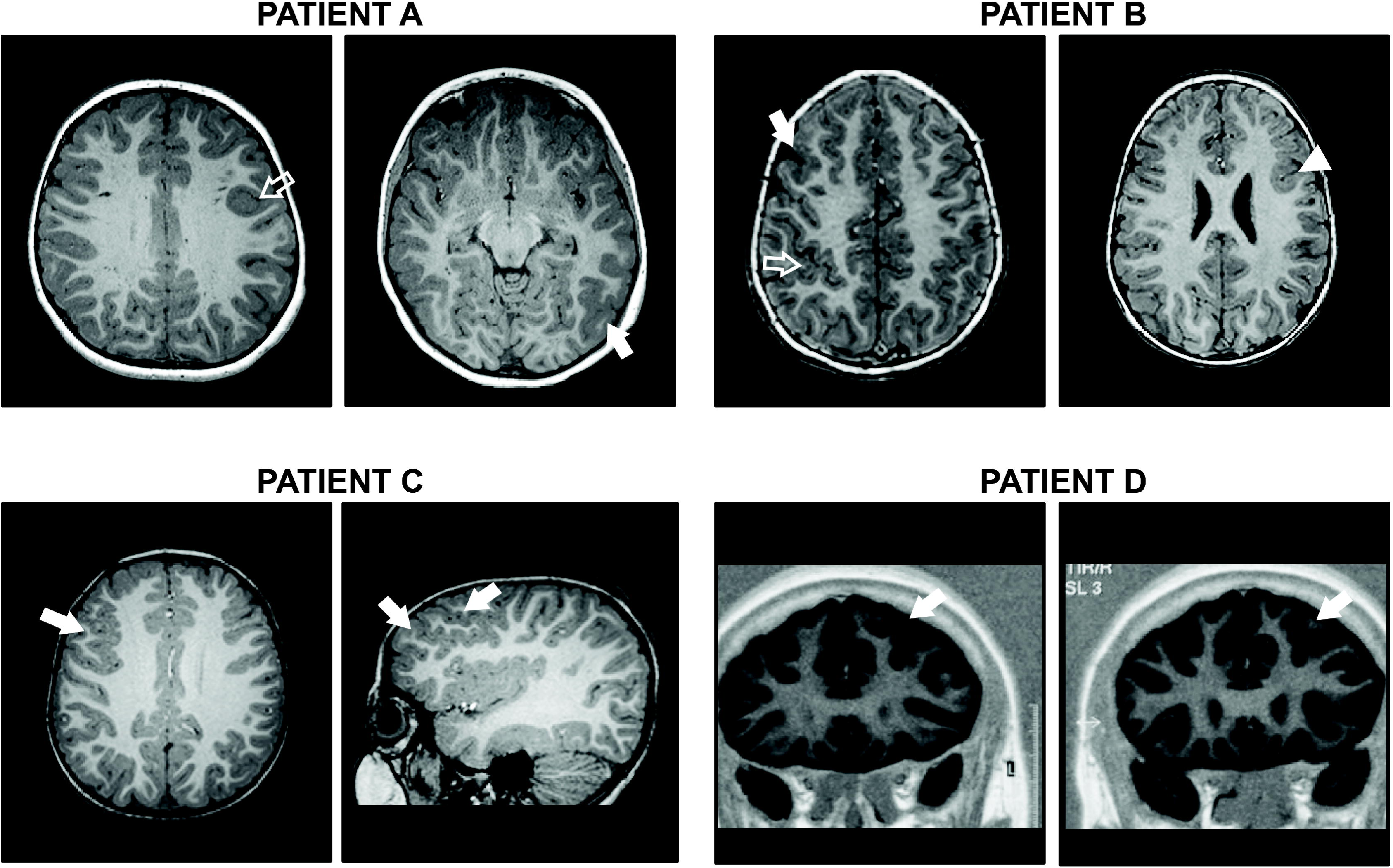
Mosaic expression of *PCDH19* results in variable cortical folding abnormalities in *PCDH19* epilepsy patients. Patient A: A focal area of cortical thickening in the left mid frontal lobe (open arrow) and mild cortical thickening and blurring of the grey/white interface in the left posterior temporal lobe was observed in axial T1 weighted MRI images (closed arrow). Patient B: Axial T1 weighted MRI images show an unusual stellate configuration to the right central sulcus with loss of cortical clarity (open arrow) and a focal retraction/puckered appearance of right lateral frontal cortex with complex sulcation and deep cortical thickening suggesting a likely bottom of the sulcus dysplasia (closed arrow). A focal area of suspected cortical dysplasia was identified in the left mid frontal region in addition to cortical thickening (arrowhead). Patient C: Axial and sagittal images show an unusual complex sulcal arrangement present in the right frontal lobe with 2 parallel sulci forming an ovoid ring configuration (closed arrows). Patient D: Contiguous coronal T1 images reveal cortical thickening with blurring of the grey/white junction which is highly suggestive of a left cortical dysplasia (closed arrows).

## Discussion

In summary, the generation of differential adhesion codes in neural progenitor cells caused by mosaic expression of *Pcdh19* appears to be the underlying cellular mechanism responsible for the unique X-linked inheritance pattern of *PCDH19*-GCE. Furthermore, our data suggests that uniform removal of PCDH19, as seen in hemizygous males, does not lead to the formation of incompatible adhesion codes allowing for normal positioning of neuroprogenitor cells and neural activity (Figure S12). The abnormal rearrangement of neuroprogenitor cells in the incipient cortex of heterozygous brains indicates that at least some of their neuronal progeny will be aberrantly positioned, regardless of whether they maintain expression of *Pcdh19* postnatally. This rearrangement has the potential to perturb functional boundaries within the cortex and alter connectivity between cortical and subcortical regions.

If the entire cortex was to undergo segregation it is likely this would result in abnormal organisation of functional cortical columns. Interestingly, cortical column organisation has been reported in gyrencephalic (folded) brains such as apes, monkeys and ferrets but there is little evidence supporting cortical column organisation in the lissencephalic (smooth) brains of mice and rats (Chow et al., 1971; Shaw et al., 1975; Tiao and Blakemore, 1976). If abnormal cell sorting does occur throughout the entire cortex this difference in neuronal architecture between humans and mice may contribute to the more severe phenotype seen in *PCDH19*-GCE patients. Furthermore, the identification of abnormal cortical sulcation in multiple *PCDH19*-GCE patients, suggests that aberrant cell sorting may generate different morphological phenotypes in gyrencephalic brains compared to lissencephalic brains. The cortical folding of human brains is largely caused by variable proliferation of basal radial glia, a cell type which represents only a minute proportion of progenitor cells in lissencephalic brains when compared to gyrencephalic brains (Fietz et al., 2010; Wang et al., 2011). Proliferation of basal radial glia cause “wedges” of cell dense areas, which ultimately cause the brain to fold. It is possible that if segregation occurred within the basal radial glial cells in the developing human brain then abnormal folding may occur due to irregular formation of “wedges”. These hypothesises could be investigated by using CRISPR-Cas9 technology to generate a *Pcdh19^+/-^* ferret (Kou et al., 2015), a gyrencephalic animal model.

More broadly, our data suggests that neurodevelopmental disorders associated with mutation of other NC PCDH family members (Bray et al., 2002; Ishizuka et al., 2016; Morrow et al., 2008; 2014) may be caused by disruption of adhesion codes. Since all other NC PCDHs are autosomal, mosaic disruption of adhesion codes would have to occur through a process other than X-inactivation. There is strong evidence that NC PCDHs are subject to random monoallelic expression (RMAE) (Savova et al., 2015). Individuals with a heterozygous germ line mutation in a given autosomal NC PCDH would have a proportion of their cells expressing either the functional or non-functional allele, resulting in disrupted adhesion codes. The proportion of RMAE affected cells would likely impact the penetrance or expressivity of any resultant phenotype.

Although both clustered and NC PCDHs can function in combinatorial adhesion complexes, our data indicates that the interaction of matching adhesion codes for each of these closely-related protein families *in vivo* can lead to different outcomes. Matching codes of clustered PCDHs are thought to result in repulsion which has been implicated in neuronal self-avoidance (Rubinstein et al., 2015; Schreiner and Weiner, 2010; Thu et al., 2014). In contrast, our data indicate that cells expressing matching NC PCDH codes selectively associate *in vivo*. Given the complex and overlapping expression pattern of NC PCDHs throughout development it seems likely that this property may direct spatial positioning of neuroprogenitors and could conceivably be utilized during morphogenesis of other organs.

The ability of cells with different identities to self-associate and form discrete populations was first identified in the early 1900s by mixing sponge cells from different colored species (Galtsoff, 1923, 1925; Wilson, 1907). We now appreciate the critical role of adhesion molecules in regulating cell identity and directing tissue morphogenesis in many developmental contexts including in the brain (Bello et al., 2012; Duguay et al., 2003; Foty and Steinberg, 2005; Price et al., 2002). Our findings advance this field by providing evidence that perturbation of cellular adhesion codes underlies the unique X-linked inheritance pattern of *PCDH19*-GCE and suggests that just a single difference in PCDH expression is enough to disrupt the complex adhesion codes present within the developing mammalian cortex.

## Acknowledgments

We thank Dr L. Luo (Stanford University) and members of the Thomas lab for discussion and comments on the manuscript. We thank the patients and their families for participating in our research, together with their referring clinicians. D.T.P was supported by an Australian Government Research Training Program Scholarship. This work was supported by an NHMRC Program Grant (400121) and the PCDH19 Alliance. The supplementary materials contain additional data. Author contributions: D. T. P., J.G., J. N. H. and P. Q. T. conceived and designed the study; D.T.P. performed and analysed all experiments except the ECoG recordings which were performed by K. L. R. and S. P; S. G. P. performed the microinjection of mouse zygotes; recruitment and phenotyping of the patient cohort was performed by R.C.D. and I. E. S. and S. A. M. analysed the MRI images.; D. T. P. prepared all figures; D. T. P., J. N. H., and P. Q. T. wrote the manuscript; and all authors revised the manuscript.

## References

Bello, S.M., Millo, H., Rajebhosale, M., and Price, S.R. (2012). Catenin-dependent cadherin function drives divisional segregation of spinal motor neurons. J. Neurosci. Off. J. Soc. Neurosci. 32, 490–505.

Bray, N.J., Kirov, G., Owen, R.J., Jacobsen, N.J., Georgieva, L., Williams, H.J., Norton, N., Spurlock, G., Jones, S., Zammit, S., et al. (2002). Screening the human protocadherin 8 (PCDH8) gene in schizophrenia. Genes Brain Behav. 1, 187–191.

Chow, K.L., Masland, R.H., and Stewart, D.L. (1971). Receptive field characteristics of striate cortical neurons in the rabbit. Brain Res. 33, 337–352.

Cooper, S.R., Jontes, J.D., and Sotomayor, M. (2016). Structural determinants of adhesion by Protocadherin-19 and implications for its role in epilepsy. eLife 5.

Depienne, C., Bouteiller, D., Keren, B., Cheuret, E., Poirier, K., Trouillard, O., Benyahia, B., Quelin, C., Carpentier, W., Julia, S., et al. (2009). Sporadic Infantile Epileptic Encephalopathy Caused by Mutations in PCDH19 Resembles Dravet Syndrome but Mainly Affects Females. PLoS Genet 5, e1000381.

Dibbens, L.M., Tarpey, P.S., Hynes, K., Bayly, M.A., Scheffer, I.E., Smith, R., Bomar, J., Sutton, E., Vandeleur, L., Shoubridge, C., et al. (2008). X-linked protocadherin 19 mutations cause female-limited epilepsy and cognitive impairment. Nat. Genet. 40, 776–781.

Duguay, D., Foty, R.A., and Steinberg, M.S. (2003). Cadherin-mediated cell adhesion and tissue segregation: qualitative and quantitative determinants. Dev. Biol. 253, 309–323.

Duszyc, K., Terczynska, I., and Hoffman-Zacharska, D. (2014). Epilepsy and mental retardation restricted to females: X-linked epileptic infantile encephalopathy of unusual inheritance. J. Appl. Genet.

Fietz, S.A., Kelava, I., Vogt, J., Wilsch-Bräuninger, M., Stenzel, D., Fish, J.L., Corbeil, D., Riehn, A., Distler, W., Nitsch, R., et al. (2010). OSVZ progenitors of human and ferret neocortex are epithelial-like and expand by integrin signaling. Nat. Neurosci. 13, 690–699.

Foty, R.A., and Steinberg, M.S. (2005). The differential adhesion hypothesis: a direct evaluation. Dev. Biol. 278, 255–263.

Gaitan, Y., and Bouchard, M. (2006). Expression of the δ-protocadherin gene Pcdh19 in the developing mouse embryo. Gene Expr. Patterns 6, 893–899.

Galtsoff, P.S. (1923). The awœboid movement of dissociated sponge cells. Biol. Bull. 45, 153–161.

Galtsoff, P.S. (1925). Regeneration after dissociation (an experimental study on sponges) I. Behavior of dissociated cells of microciona prolifera under normal and altered conditions. J. Exp. Zool. 42, 183–221.

Garrett, A.M., and Weiner, J.A. (2009). Control of CNS synapse development by ?-protocadherin-mediated astrocyte-neuron contact. J. Neurosci. Off. J. Soc. Neurosci. 29, 11723–11731.

Guerrini, R., Marini, C., and Mantegazza, M. (2014). Genetic epilepsy syndromes without structural brain abnormalities: clinical features and experimental models. Neurother. J. Am. Soc. Exp. Neurother. 11, 269–285.

Higurashi, N., Nakamura, M., Sugai, M., Ohfu, M., Sakauchi, M., Sugawara, Y., Nakamura, K., Kato, M., Usui, D., Mogami, Y., et al. (2013). PCDH19-related female-limited epilepsy: further details regarding early clinical features and therapeutic efficacy. Epilepsy Res. 106, 191–199.

International League Against Epilepsy Consortium on Complex Epilepsies. Electronic address: (2014). Genetic determinants of common epilepsies: a meta-analysis of genome-wide association studies. Lancet Neurol. 13, 893–903.

Ishizuka, K., Kimura, H., Wang, C., Xing, J., Kushima, I., Arioka, Y., Oya-Ito, T., Uno, Y., Okada, T., Mori, D., et al. (2016). Investigation of Rare Single-Nucleotide PCDH15 Variants in Schizophrenia and Autism Spectrum Disorders. PLoS ONE 11, e0153224.

Kim, S.-Y., Chung, H.S., Sun, W., and Kim, H. (2007). Spatiotemporal expression pattern of non-clustered protocadherin family members in the developing rat brain. Neuroscience 147, 996–1021.

Kou, Z., Wu, Q., Kou, X., Yin, C., Wang, H., Zuo, Z., Zhuo, Y., Chen, A., Gao, S., and Wang, X. (2015). CRISPR/Cas9-mediated genome engineering of the ferret. Cell Res. 25, 1372–1375.

Krishna-K, C.R. (2009). Expression of cadherin superfamily genes in brain vascular development. J. Cereb. Blood Flow Metab. 29, 224–229.

Lefebvre, J.L., Kostadinov, D., Chen, W.V., Maniatis, T., and Sanes, J.R. (2012). Protocadherins mediate dendritic self-avoidance in the mammalian nervous system. Nature 488, 517–521.

Miller, J.A., Ding, S.-L., Sunkin, S.M., Smith, K.A., Ng, L., Szafer, A., Ebbert, A., Riley, Z.L., Royall, J.J., Aiona, K., et al. (2014). Transcriptional landscape of the prenatal human brain. Nature 508, 199–206.

Morrow, E.M., Yoo, S.-Y., Flavell, S.W., Kim, T.-K., Lin, Y., Hill, R.S., Mukaddes, N.M., Balkhy, S., Gascon, G., Hashmi, A., et al. (2008). Identifying autism loci and genes by tracing recent shared ancestry. Science 321, 218–223.

Ozawa, M., and Kemler, R. (1998). Altered cell adhesion activity by pervanadate due to the dissociation of alpha-catenin from the E-cadherin.catenin complex. J. Biol. Chem. 273, 6166–6170.

Pederick, D.T., Homan, C.C., Jaehne, E.J., Piltz, S.G., Haines, B.P., Baune, B.T., Jolly, L.A., Hughes, J.N., Gecz, J., and Thomas, P.Q. (2016). Pcdh19 Loss-of-Function Increases Neuronal Migration In Vitro but is Dispensable for Brain Development in Mice. Sci. Rep. 6, 26765.

Price, S.R., De Marco Garcia, N.V., Ranscht, B., and Jessell, T.M. (2002). Regulation of Motor Neuron Pool Sorting by Differential Expression of Type II Cadherins. Cell 109, 205–216.

Rubinstein, R., Thu, C.A., Goodman, K.M., Wolcott, H.N., Bahna, F., Mannepalli, S., Ahlsen, G., Chevee, M., Halim, A., Clausen, H., et al. (2015). Molecular Logic of Neuronal Self-Recognition through Protocadherin Domain Interactions. Cell 163, 629–642.

Ryan, S.G., Chance, P.F., Zou, C.H., Spinner, N.B., Golden, J.A., and Smietana, S. (1997). Epilepsy and mental retardation limited to females: an X-linked dominant disorder with male sparing. Nat. Genet. 17, 92–95.

Savova, V., Patsenker, J., Vigneau, S., and Gimelbrant, A.A. (2015). dbMAE: the database of autosomal monoallelic expression. Nucleic Acids Res. gkv1106.

Scheffer, I.E., Turner, S.J., Dibbens, L.M., Bayly, M.A., Friend, K., Hodgson, B., Burrows, L., Shaw, M., Wei, C., Ullmann, R., et al. (2008). Epilepsy and mental retardation limited to females: an under-recognized disorder. Brain 131, 918–927.

Schreiner, D., and Weiner, J.A. (2010). Combinatorial homophilic interaction between gamma-protocadherin multimers greatly expands the molecular diversity of cell adhesion. Proc. Natl. Acad. Sci. U. S. A. 107, 14893–14898.

Semple, B.D., Blomgren, K., Gimlin, K., Ferriero, D.M., and Noble-Haeusslein, L.J. (2013). Brain development in rodents and humans: Identifying benchmarks of maturation and vulnerability to injury across species. Prog. Neurobiol. 0, 1–16.

Shaw, C., Yinon, U., and Auerbach, E. (1975). Receptive fields and response properties of neurons in the rat visual cortex. Vision Res. 15, 203–208.

Sun, T., and Hevner, R.F. (2014). Growth and folding of the mammalian cerebral cortex: from molecules to malformations. Nat. Rev. Neurosci. 15, 217–232.

Tai, K., Kubota, M., Shiono, K., Tokutsu, H., and Suzuki, S.T. (2010). Adhesion properties and retinofugal expression of chicken protocadherin-19. Brain Res. 1344, 13–24.

Thu, C.A., Chen, W.V., Rubinstein, R., Chevee, M., Wolcott, H.N., Felsovalyi, K.O., Tapia, J.C., Shapiro, L., Honig, B., and Maniatis, T. (2014). Generation of single cell identity by homophilic interactions between combinations of α, β and ? protocadherins. Cell 158, 1045–1059.

Tiao, Y.C., and Blakemore, C. (1976). Functional organization in the visual cortex of the golden hamster. J. Comp. Neurol. 168, 459–481.

Uemura, M., Nakao, S., Suzuki, S.T., Takeichi, M., and Hirano, S. (2007). OL-protocadherin is essential for growth of striatal axons and thalamocortical projections. Nat. Neurosci. 10, 1151–1159.

Wang, X., Tsai, J.-W., LaMonica, B., and Kriegstein, A.R. (2011). A new subtype of progenitor cell in the mouse embryonic neocortex. Nat. Neurosci. 14, 555–561.

Wilson, H.V. (1907). On some phenomena of coalescence and regeneration in sponges. J. Exp. Zool. 5, 245–258.

